# The Limits of Sequence-Based Biosecurity Screening Tools in the Age of AI-Assisted Protein Design

**DOI:** 10.64898/2026.03.04.709671

**Authors:** Bruce J. Wittmann, Nicole E. Wheeler, Steven T. Murphy, Tom Mitchell, Brittany Magalis, Bryan T. Gemler, Kevin Flyangolts, James Diggans, Adam Clore, Jacob Beal, Craig Bartling, Tessa Alexanian, Eric Horvitz

## Abstract

Rapid advancements in AI have enabled significant progress in protein and nucleic acid design, but they also pose biosecurity challenges. We examine the vulnerabilities of biosecurity screening software (BSS) to AI-reformulated synthetic homologs of proteins of concern (POCs) that have been fragmented into smaller segments. We evaluate four BSS tools that were recently patched to enhance their AI resiliency. Without any further modification, we found that two of the four tools were capable of robustly detecting fragments as short as 50 nucleotides, demonstrating screening capabilities that exceed those requested in the U.S. Framework for Nucleic Acid Synthesis. Upgraded versions of the other two tools improved performance. Although our findings confirm the effectiveness of the tested BSS tools, at the same time, they emphasize the urgency of developing alternate BSS approaches to counter evolving AI-enabled biosecurity risks.

---

The rapid advancement of AI methods has revolutionized biotechnology, dramatically expanding capabilities of protein and nucleic acid design.^1–6^ However, these innovations can also introduce new biosecurity risks.^7–15^ Recently, we demonstrated a vulnerability where AI-assisted protein design (AIPD) tools could be used to evade the biosecurity screening software (BSS) used by nucleic acid synthesis providers to screen customer orders for sequences of concern (SOCs).^16^ Specifically, we showed how such tools could be used to reformulate wild-type proteins of concern (POCs) to produce *synthetic homologs* with minimal sequence identity to those wild types but that could nevertheless fold into structurally similar proteins, and thus potentially retain their function. Because BSS tools used by most nucleic acid synthesis providers rely on sequence identity to known SOCs to flag potential hazards,^17–20^ these sequence-divergent synthetic homologs could be difficult to detect, potentially allowing malicious actors to obtain proteins with hazardous functions (e.g., toxins or viral proteins).

Importantly, we showed that those tools could be updated to reduce their vulnerability to synthetic homologs of POCs. Further, a follow-up wet-lab study has provided additional reassurance by investigating the efficacy of an end-to-end synthetic homolog pipeline with benign proteins, providing evidence that tested AIPD tools may not be effective enough to produce proteins that both maintain function and are sufficiently sequence-divergent to evade sequence-based BSS tools updated to be AI resilient.^21^ Nonetheless, concerns persist about the potential future capabilities of AIPD tools, and we can expect the effectiveness of sequence-based BSS to decline with continuing improvement of AIPD capabilities. Thus, we must work to develop alternate BSS tools where SOCs are more directly identified (e.g., with methods that could predict a sequence’s function) rather than via a correlate like sequence identity. Simultaneously, though, we must also determine the limits of current sequence-centric BSS tools to gauge the urgency with which these alternate detection strategies must be pursued.

In this study, we test the limits of current BSS tools by evaluating their ability to identify *fragments* of synthetic homologs. Specifically, we evaluate the abilities of the screening tools to identify fragments of synthetic homologs whose full-length sequences were appropriately flagged as hazardous in our earlier study (Fig. 1, *Methods: Fragmentation*). This study is motivated by concerns about the possibility that, rather than ordering full-length genes, malevolent actors might instead order fragments thereof that could be assembled downstream, and that those fragments would be more difficult for BSS systems to detect.

**Fig. 1.**
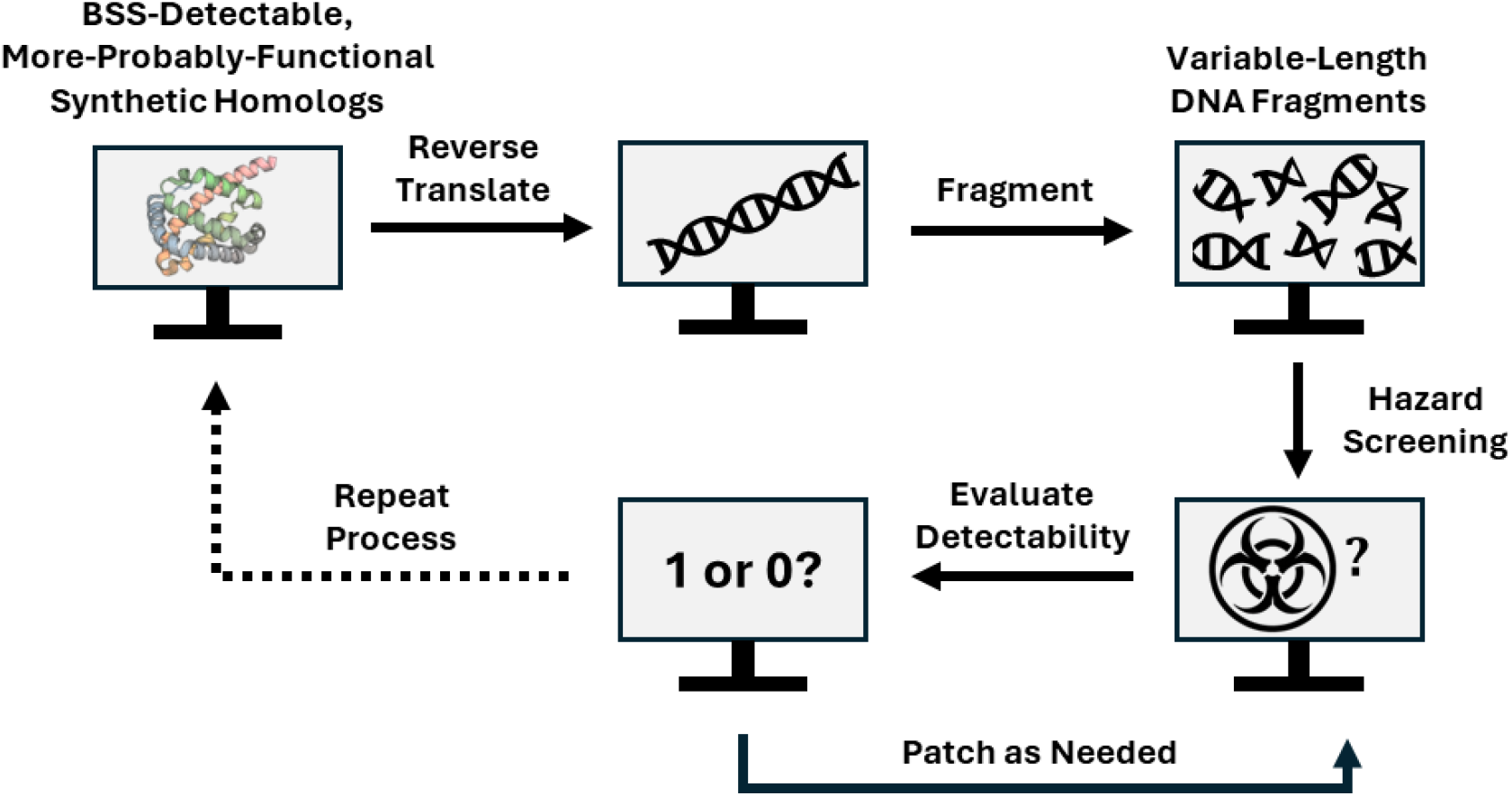
Summary of workflow.

We start with an exploration of the performance of BSS tools on fragment lengths of 200 nucleotides (66 amino acids) per specification in the 2024 Framework for Nucleic Acid Synthesis issued by the Office of Science and Technology Policy (OSTP) that screening tools should identify fragments of SOCs coded by 200 nucleotides or longer.^22^ We then explore screening capabilities on progressively shorter fragments, decreasing the length in 25-nucleotide increments down to a minimum of 25 nucleotides (7 or 8 amino acids, depending on the reading frame). The lower limit of 25 nucleotides was included to test BSS tools beyond the boundary of the 50-nucleotide fragment length that OSTP recommends BSS systems should be capable of handling by October 2026. It is worth noting that all BSS tools tested in this study have demonstrated their capabilities of detecting 50-nucleotide-long or shorter fragments of wild-type SOCs;^23^ this study tests whether those capabilities generalize to synthetic homologs.

A summary of results derived from passing fragmented true-positive (i.e., synthetic homologs of SOCs) and true-negative (i.e., synthetic homologs of benign proteins) sequences through the final versions of tools studied in our original work is presented in Fig. 2. The chart displays performance in terms of Matthew’s correlation coefficient (MCC, *Methods*: *Matthew’s Correlation Coefficient*) for each provider and tool at each fragment length, where results are reported as “Provider N, Tool X.” This metric is a classification analog to the more familiar Pearson and Spearman correlation coefficients. Like them, it ranges between −1 and 1, with a negative score indicating inverse correlation (i.e., the wrong label is assigned more frequently, corresponding to high false positive and/or false negative rates) and a positive score indicating positive correlation (i.e., the correct label is assigned more frequently, corresponding to high true positive and/or true negative rates). An effective BSS tool should thus have a high, positive MCC.

**Fig. 2.**
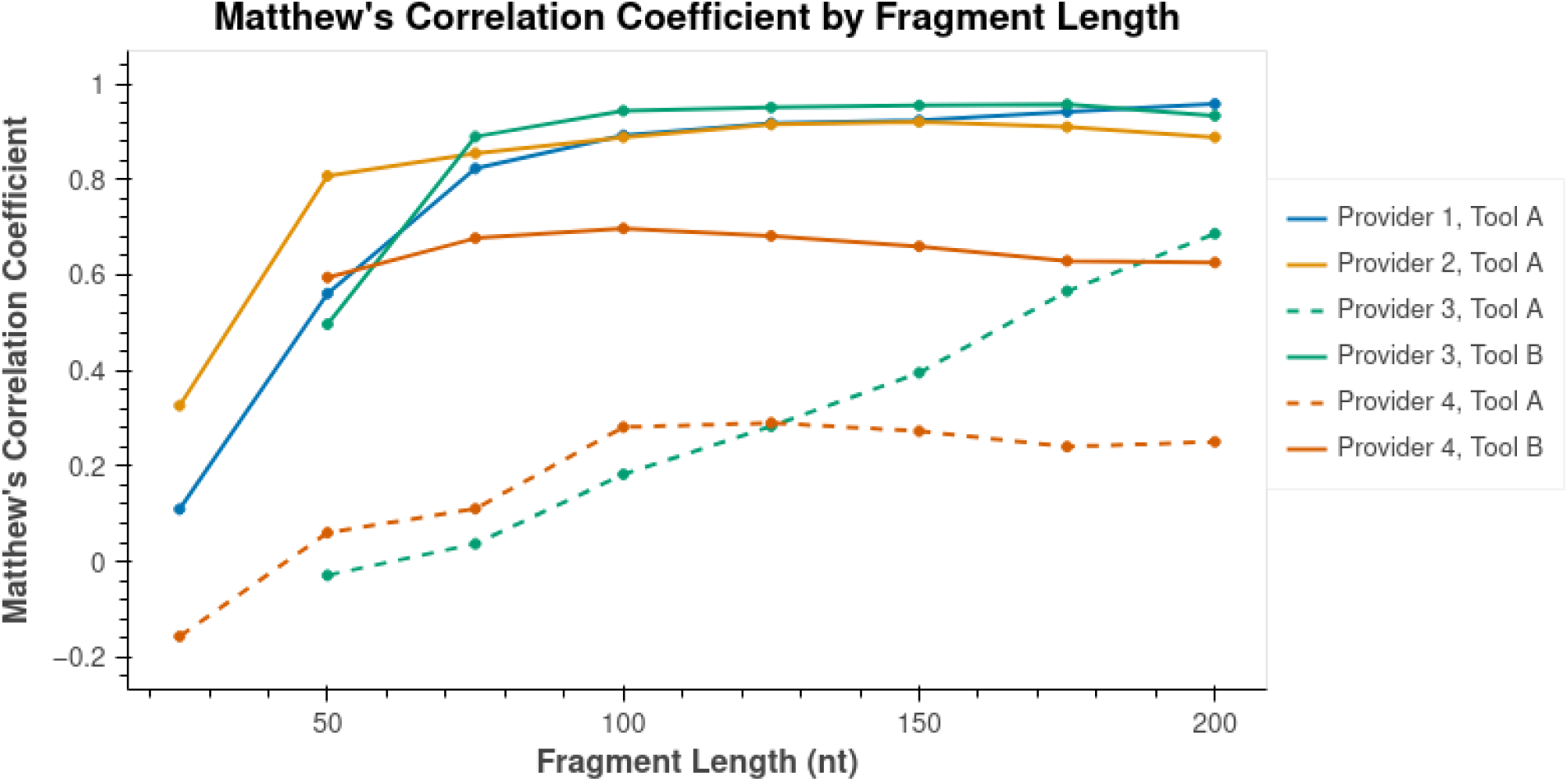
Sequence-level analysis. The x-axis represents fragment length while the y-axis gives Matthew’s correlation coefficient (MCC). Some tools lack a datapoint at 25 nt as no sequences were flagged regardless of ground truth, leading to an undefined MCC. Note that “Tool A” is used to designate original tools from given providers while “Tool B” is used to designate the updated ones. See also, Fig. S1–S6, Panels 1.

Considering a given synthetic homolog to be flagged if at least one fragment derived from that sequence was flagged (hereafter, “sequence-level analysis”), we found that the performance of Providers 1 and 2’s Tools A on fragments was consistent with their performance on full-length synthetic homologs down to approximately the 50-nucleotide range, with both tools exhibiting high MCC, true negative, and true positive rates (Fig. 2, Table S1, Data S1). In contrast, Provider 3’s Tool A struggled to detect true-positive fragments, particularly as fragment length declined, thus lowering its MCC (Table S1). Provider 4’s Tool A showed the opposite problem, with its MCC being lowered across all fragment sizes due to a high false positive rate (Table S1). Alternate tools from these two providers (“Tools B”), however, which were developed either prior to these initial results or in in response to them, yielded improved results (Table S1).

In an applied setting where BSS tools are deployed to screen live customer orders, the sequence-level analysis described in the previous paragraph—where an order is flagged if any sequence it contains is flagged—is sensible from a security perspective, as a single harmful fragment can indicate malicious intent, and so should trigger analysis of an order as a whole. However, flagging an order if any sequence it contains is flagged is not as helpful for answering the central motivation of this follow-up work, which was to determine the current limit of sequence-based BSS tools. To better answer this question, we also evaluated tool performance at the *fragment level* as a function of both the sequence identity and *longest common subsequence* (LCS) shared between synthetic homolog fragments and their analogous wild-type fragments (hereafter, “fragment-level analysis”).

Fig. 3 shows the results of this fragment-level analysis using the most up-to-date BSS tools, demonstrating trends that we expect for sequence-based screening strategies: As sequence identity and LCS drop, the rate at which fragments were flagged drops as well. Interestingly, different tools have different boundaries at which flagging rates begin to decline. Provider 1’s tool, for example, flagged nearly 100% of fragments with an LCS greater than 20, with performance dropping below this threshold. Sequence identity appears to have limited effect on flagging rates until the ~30% identity mark is reached, a point at which all fragments already have an LCS < 20, and beyond which flagging rates dropped to near 0%. The performances of the other three providers’ tools, by contrast, exhibited greater covariance with the two sequence metrics. Provider 2’s tool’s performance, for example, only dropped off when both LCS was below ~10 *and* percent identity was below 50; Providers 3 and 4’s Tools B show a similar multivariate-dependent boundary.

**Fig. 3.**
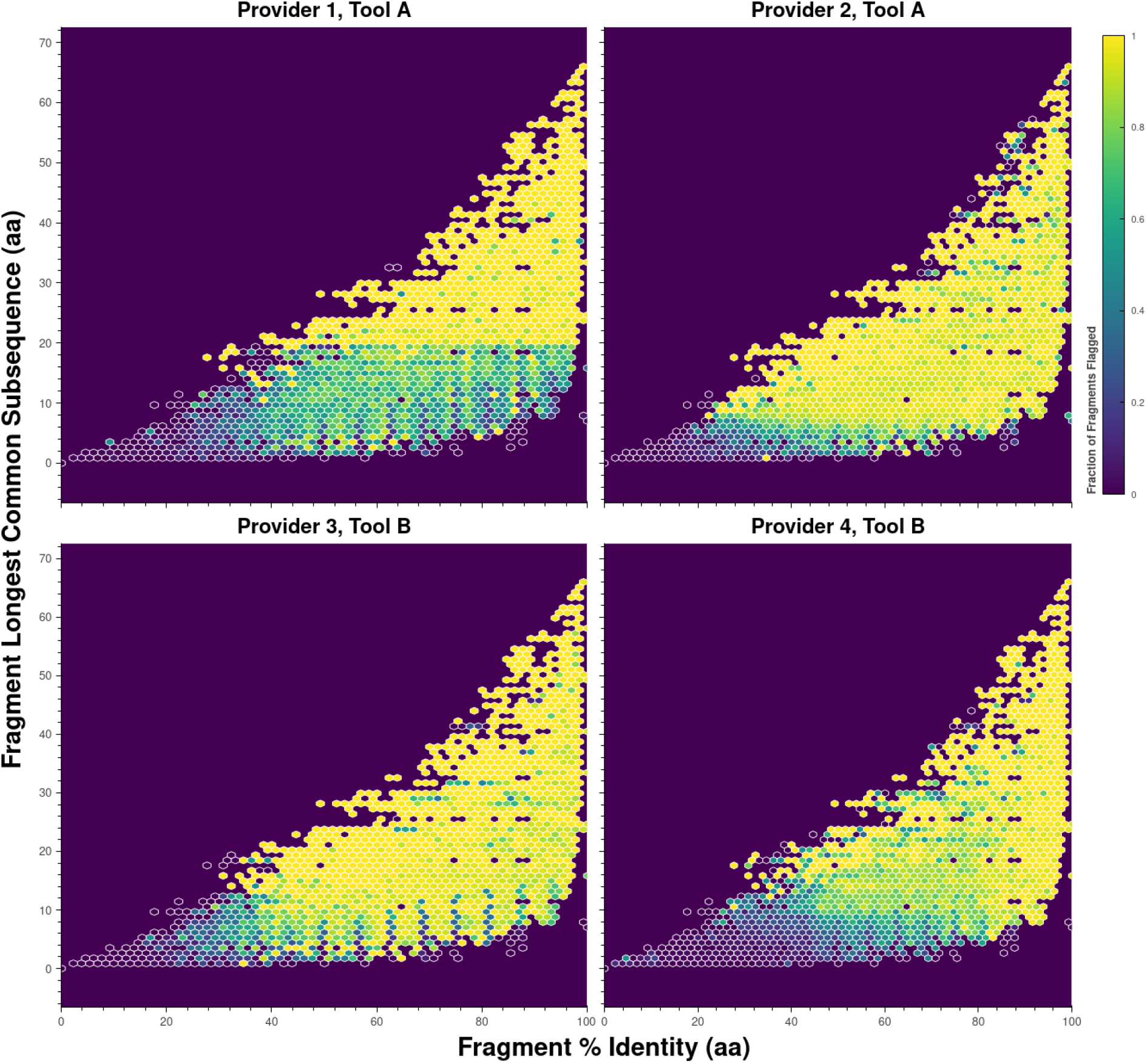
Fragment-level analysis. For each subplot, the x-axis displays the percent identity of a fragment relative to its analogous wild-type fragment. The y-axis gives the longest common subsequence (LCS) shared by those two fragments. Each hexagon is colored according to the rate with which the fragments it contains were flagged. See also, Fig. S1–S6, Panels 2–3.

Both the sequence- and fragment-level results strengthen confidence in the robustness of patched BSS tools against AI-reformulated SOCs. For all tools tested, even a determined adversary—one who reformulates and fragments protein sequences in an attempt to evade detection—is still likely to have orders flagged by the BSS tools tested here. *When combined with the wet-laboratory results discussed earlier*,^*21*^ *these findings suggest that sequence-based BSS tools patched for AI-resilience are effective against widely used AIPD capabilities*.

Despite these findings, current technology has clear limitations. As evidenced by performance drop-offs below certain LCS and sequence identity thresholds (Fig. 3), if AIPD tools are developed that can be used to design functional sequences with minimal similarity to wild-type proteins, existing BSS tools will be rendered ineffective. Further, the relatively wide array of sequence-level performance measured across the four tools for 50-nucleotide fragments (ranging from ~0.5 to 0.8 MCC, Fig. 2) represents an additional risk for nucleic acid synthesis providers selecting tools to secure their order streams: A tool that performs well against traditional threats may be less effective against highly engineered threats, and it is not clear how a provider might prioritize defending against one over the other, per preferences and risk profiles over the identified tradeoffs.

A critical unknown remains: Given 50-nucleotide fragments alone as a signal, we do not know if it is possible to predict whether a highly engineered synthetic homolog (i.e., one with even lower sequence-identity-to-wild-type than the synthetic homologs tested here) or truly de novo-designed protein sequence encodes a function of concern. As fragment size decreases, the probability that a fragment is unique to a threat also decreases, necessitating strategies beyond sequence-based screening alone. Longer engineered sequences can be folded in silico, allowing structural similarities to known SOCs to be calculated and used as higher-resolution proxies for function detection than sequence similarity. However, this technique will not work for shorter oligonucleotide sequences in isolation. For those, direct prediction of sequence function may be necessary, though exactly how feasible this would be—particularly when analyzing highly mutated fragments—remains unclear. It is likely that broader context, such as evidence provided by other sequences in the same or other orders as well as customer information, would be required to make a reliable determination of threat.

While early efforts to develop such function-prediction and contextually aware BSS have begun, none are currently operationally deployable. Fortunately, current AIPD models appear to be largely incapable of producing the extremely divergent (<30% sequence identity) sequences that would necessitate adoption of such alternate strategies. It is therefore imperative to proceed with caution, maintain vigilant oversight of AIPD advancements, continually reassess AIPD tool capabilities against current BSS benchmarks, and continue to act with urgency to create BSS tools that can meet tomorrow’s threats.

## Methods

### Fragmentation

Fragments were derived from a subset of sequences generated as part of our earlier efforts to patch BSS tools to be AI-resilient.^16^ The experimental set of fragments was built via the following procedure:

1. From the full set of sequences generated as part of our original work, non-wild-type sequences with a TM Score > 0.5 and ΔpLDDT > −10.0— those termed “more-probably-functional” in that work—were selected.
2. The subset resulting from Step 1 was further filtered to include only sequences flagged by at least three of the four screening tools in the original study.
3. Ten sequences were randomly sampled without replacement from each group of sequences generated using the same tool, template protein, and conserved residues (see details on the generation procedure in the original study for more details on these different groups). If fewer than ten sequences met the criteria, all sequences for that group were included.
4. The sets sampled in Step 3 and wild-type template sequences were combined to build a final set of 5505 more-probably-functional SOCs. The flagging rates of the subsampled sequences were reflective of the broader population of more-probably-functional sequences reported in our original study (Table S2, see Table S3 of Wittmann et al, 2025).^16^
5. DNA sequences encoding each of the 5505 proteins were fragmented into lengths of 25, 50, 75, 100, 125, 150, 175, and 200 nucleotides, with overlap lengths of 5 nucleotides for fragments 25 nucleotides long and overlap lengths of 25 nucleotides for all longer fragments. If the last fragment in a DNA sequence was shorter than the target, then that fragment was discarded. In total, this procedure resulted in ~700k fragments.

To build the “true negative” set of fragments, Steps 3–5 were repeated for the “Conditionally Generated True Negatives” and “Known True Negatives” sets from our original study (see “Negative Controls,” Items 2 and 3 in Wittmann et al., 2025 for a description of these sets).^16^ This procedure yielded 1036 negative control sequences (980 “Conditionally Generated True Negatives” and 56 “Known True Negatives”) and ~170k fragments. The fragments, their relevant statistics, and how each tool assigned flags can be found in Data S2 and Data S3 for the experimental and negative control sets, respectively.

### Biosecurity Screening Software Analysis

For each fragment length, one FASTA file was created for each of the full-length sequences with each file containing the fragments for that sequence at that length (e.g., for the experimental set, resulting in a total of 5505 sequences * 8 fragment lengths = 44,040 files). Each file was then submitted as a single order to each of the tested BSS tools, each of which returned a binary label of “flag” or “no flag” for each fragment. As described above, we considered a sequence to have been flagged if at least one fragment of a given length derived from that sequence was flagged. Details on the strategies employed by each tool can be found in our prior work.^16^

### Matthew’s Correlation Coefficient

If we defined tp as the number of true positives, tn as the number of true negatives, fp as the number of false positives, and fn as the number of false negatives, then Matthew’s Correlation Coefficient is defined as

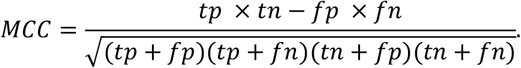

In the ideal case, where fp = fn = 0, this reduces to

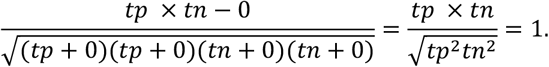

In the worst case, where tp = tn = 0, this reduces to

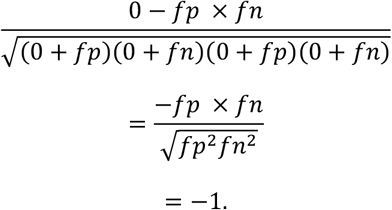

For intermediate cases, the sign of metric is determined by the relative balance of true positives and negatives to false positives and negatives. If true positives and negatives outweigh the false positives and negatives, then the sign is positive, and vice versa. The degree to which the true assignments outweigh the false assignments determines the magnitude of the metric.

## Acknowledgments

This document does not contain technology or technical data controlled under either U.S. International Traffic in Arms Regulation or U.S. Export Administration Regulations.

## CRediT Taxonomy

B.J.W.: Conceptualization, Data Curation, Formal Analysis, Investigation, Methodology, Project Administration, Resources, Software, Validation, Visualization, Writing – Original Draft, Writing – Review and Editing

N.W., S.T.M., T.M., B.M., B.T.G., K.F., J.D., A.C.,J.B., C.B., T.A.: Conceptualization, Formal Analysis, Investigation, Methodology, Software, Writing – Review and Editing

E.H.: Conceptualization, Investigation, Methodology, Project Administration, Resources, Writing – Original Draft, Writing – Review and Editing

## Data and Materials Availability

The code and data are available upon request to the International Biosecurity and Biosafety Initiative for Science (IBBIS, https://ibbis.bio/).

## Supplementary Materials for

### Supplementary Data

#### Data S1. Sequence-level classification statistics by provider, fragment length, and tool

Column headers are as follows:

1. Provider: ID of the provider.
2. Fragment Length: The length of the fragment in nucleotides.
3. TPR: True positive rate 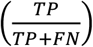
4. FPR: False positive rate 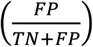
5. MCC: Matthew’s correlation coefficient. See *Methods* for definition.
6. TN: Number of true negatives.
7. TP: Number of true positives.
8. FN: Number of false negatives.
9. FP: Number of false positives.
10. Total: Sum of TN, TP, FN, and FP.
11. Tool: The ID of the tool (“A” for tools used in Wittmann et al., 2025, “B” for tools updated to be more effective at screening fragments).

#### Data S2. All data for experimental fragments

Column headers are defined as follows:

1. Fragment Index: Unique index assigned to a fragment of a specific sequence.
2. Flag: Whether this fragment was flagged as hazardous by the provider in the “Provider” column.
3. Tool: The generative model used to build the original sequence.
4. Protein: The protein used as a template. This uses the same random indexing scheme to deidentify proteins as used in Wittmann et al., 2025.
5. Condition: The strategy used to generate the protein. See “Materials and Methods: Reformulation” in the supplementary materials of Wittmann et al., 2025 for a description of the different conditions.
6. Sequence Name: The name of the sequence. This directly corresponds to the sequence name used in Wittmann et al., 2025.
7. Fragment Length: The length of the fragment in nucleotides.
8. Provider: ID of the provider/tool.
9. Upgraded: Whether or not the row corresponds to the most up-to-date BSS from the provider. This will be “TRUE” for Providers 1 and 2, Tools A, as well as for Providers 3 and 4, Tools B. It will be “FALSE” for Providers 3 and 4, Tools A.
10. DNA Fragment: The DNA sequence of the fragment.
11. Protein Fragment: The amino acid sequence of the fragment translated using the reading frame of the original, full-length sequence.
12. DNA Fragment (WT): The DNA sequence of the analogous wild-type fragment.
13. Protein Fragment (WT): The amino acid sequence of the analogous wild-type fragment.
14. Protein Frac Identity: The fractional sequence identity between the sequences in the “Protein Fragment” and “Protein Fragment (WT)” columns.
15. DNA Frac Identity: The fractional sequence identity between the sequences in the “DNA Fragment” and “DNA Fragment (WT)” columns.
16. Protein LCS: The longest common substring shared between the sequences in the “Protein Fragment” and “Protein Fragment (WT)” columns.
17. DNA LCS: The longest common substring shared between the sequences in the “DNA Fragment” and “DNA Fragment (WT)” columns.

#### Data S3. All data for negative control fragments

Column header definitions match those used for Data S2, with the following minor change:

1) Protein: Rather than a random index identifier, this is either the PDB ID or the NCBI Accession Number of the protein

## Supplementary Tables

**Table S1.** False positive rate by tool and fragment length.

| Fragment Length (nt) | Provider 1 |  | Provider 2 |  | Provider 3 |  |  |  | Provider 4 |  |  |  |
| --- | --- | --- | --- | --- | --- | --- | --- | --- | --- | --- | --- | --- |
|  | Tool A |  | Tool A |  | Tool A |  | Tool B |  | Tool A |  | Tool B |  |
|  | TPR | FPR | TPR | FPR | TPR | FPR | TPR | FPR | TPR | FPR | TPR | FPR |
| 25 | 0.0716 | 0.0000 | 0.4300 | 0.0000 | 0.0000 | 0.0000 | 0.0000 | 0.0000 | 0.0000 | 0.0290 | 0.0000 | 0.0000 |
| 50 | 0.7435 | 0.0000 | 0.9221 | 0.0000 | 0.0000 | 0.0010 | 0.6748 | 0.0010 | 0.9935 | 0.9778 | 0.8812 | 0.1988 |
| 75 | 0.9301 | 0.0010 | 0.9450 | 0.0000 | 0.0107 | 0.0010 | 0.9602 | 0.0010 | 0.9853 | 0.9411 | 0.9019 | 0.1342 |
| 100 | 0.9613 | 0.0010 | 0.9595 | 0.0000 | 0.1789 | 0.0000 | 0.9809 | 0.0000 | 0.9767 | 0.8089 | 0.9061 | 0.1158 |
| 125 | 0.9715 | 0.0010 | 0.9704 | 0.0010 | 0.3678 | 0.0087 | 0.9853 | 0.0087 | 0.9728 | 0.7896 | 0.8970 | 0.1149 |
| 150 | 0.9737 | 0.0010 | 0.9729 | 0.0029 | 0.5473 | 0.0077 | 0.9866 | 0.0077 | 0.9609 | 0.7712 | 0.8839 | 0.1139 |
| 175 | 0.9804 | 0.0010 | 0.9680 | 0.0000 | 0.7519 | 0.0048 | 0.9867 | 0.0048 | 0.9393 | 0.7500 | 0.8630 | 0.1100 |
| 200 | 0.9860 | 0.0000 | 0.9594 | 0.0000 | 0.8489 | 0.0000 | 0.9771 | 0.0000 | 0.9299 | 0.7210 | 0.8609 | 0.1110 |

**Table S2.**
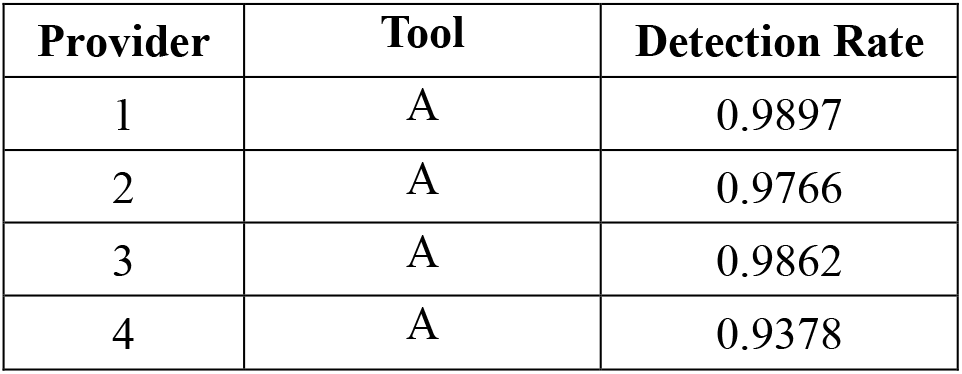
Detection rate in prior work of full-length more-probably-functional synthetic homologs of SOCs used to build fragments.^16^.

| Provider | Tool | Detection Rate |
| --- | --- | --- |
| 1 | A | 0.9897 |
| 2 | A | 0.9766 |
| 3 | A | 0.9862 |
| 4 | A | 0.9378 |

## Supplementary Figures

**Fig. S1.**
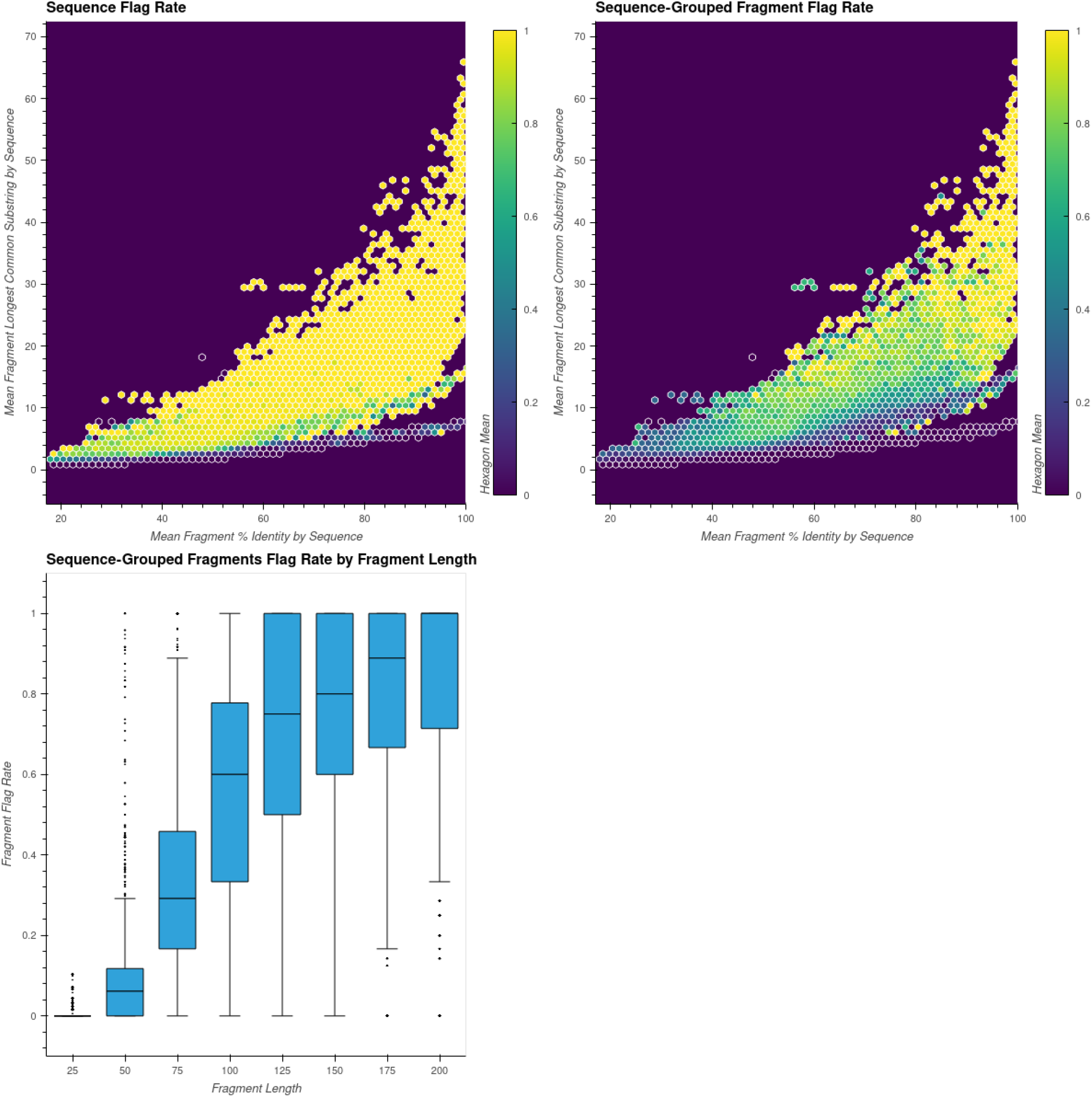
Alternate representations of data displayed in Fig. 2 and Fig. 3 for Provider 1, Tool A. The first panel (top left) shows, for a given full-length sequence, the average fragment percent identity (x-axis) against the average longest common subsequence (y-axis). Each hexagon is colored according to the rate at which *at least one fragment* derived from the sequences it contains were flagged. That is, Panel 1 is a finer-grained representation of the data in Fig. 2. The x- and y-axes of the second panel (top right) have the same meaning as the first panel, only here each hexagon is colored according to the average rate of the frequency with which fragments derived from the sequences it contains were flagged. Put differently, Panel 2 is a coarser-grained representation of the data in Fig. 3 where the mean is taken over all fragments in a sequence before the binning operation is applied to color the hexagons. The final panel (bottom left) shows the same information as the second—the fraction of fragments flagged by sequence— only now plotted against the fragment length and displayed as a box-and-whisker plot rather than binning via hexagons.

**Fig. S2.**
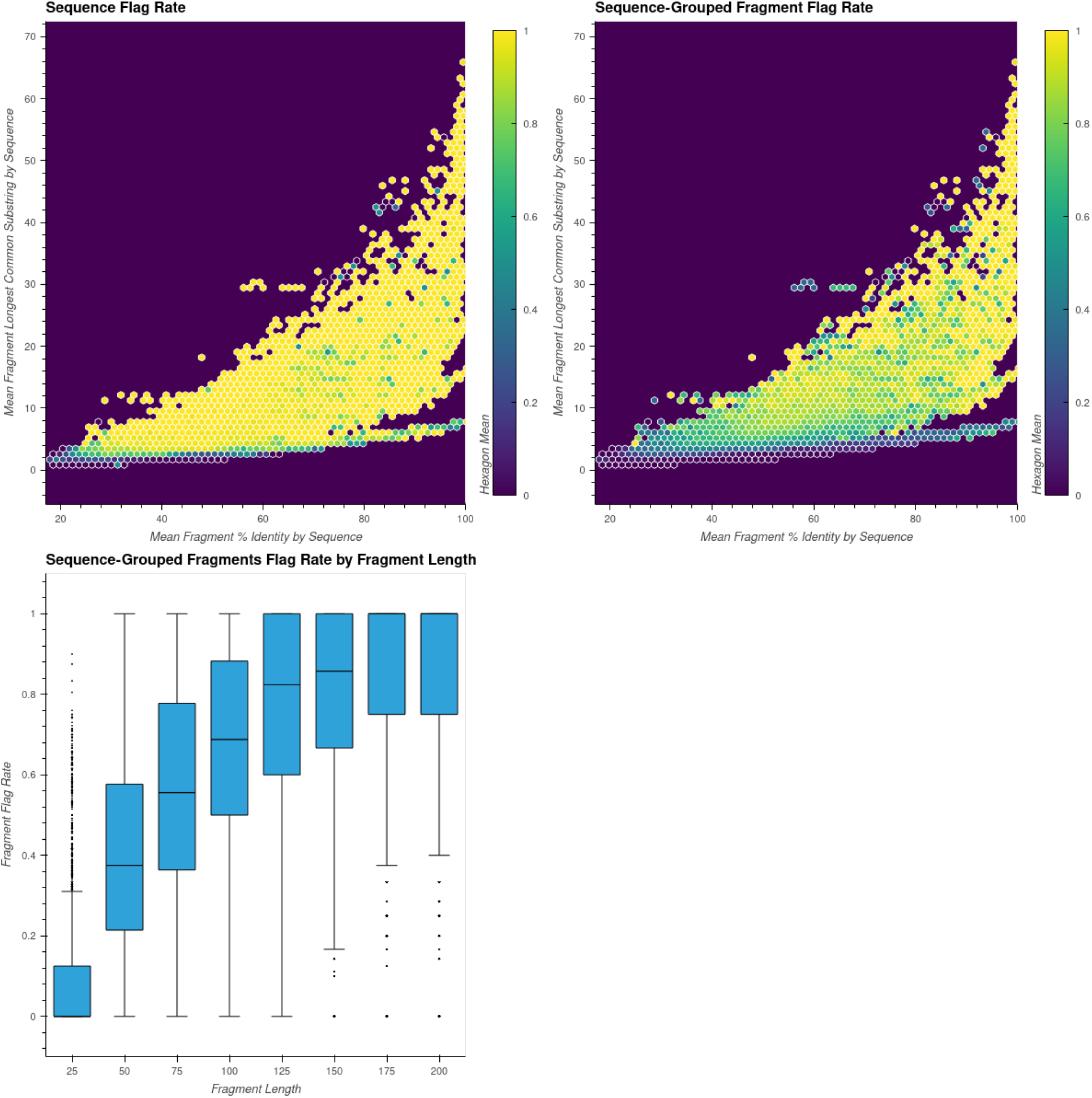
Identical to Fig. S1, but for Provider 2, Tool A.

**Fig. S3.**
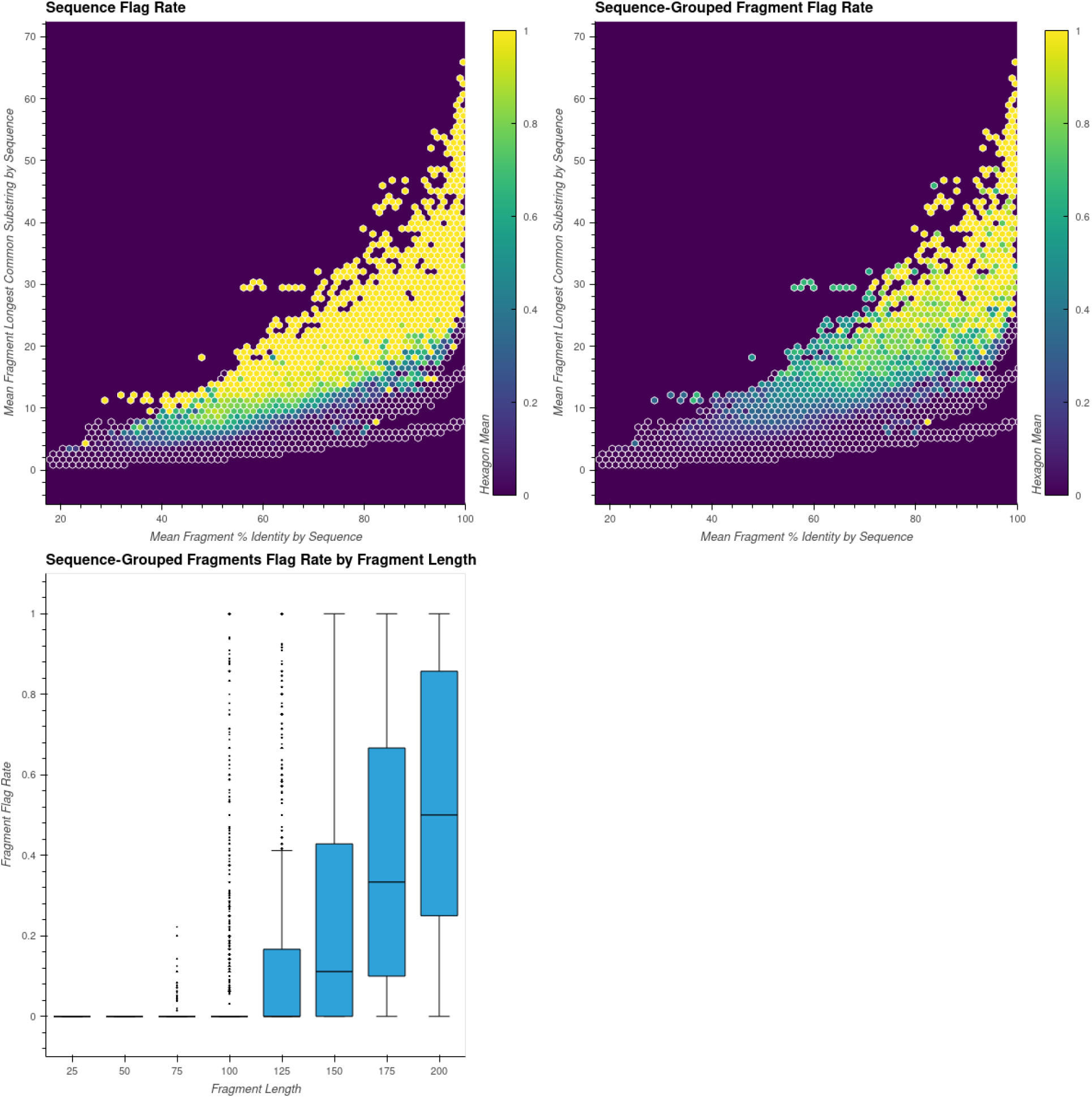
Identical to Fig. S1, but for Provider 3, Tool A.

**Fig. S4.**
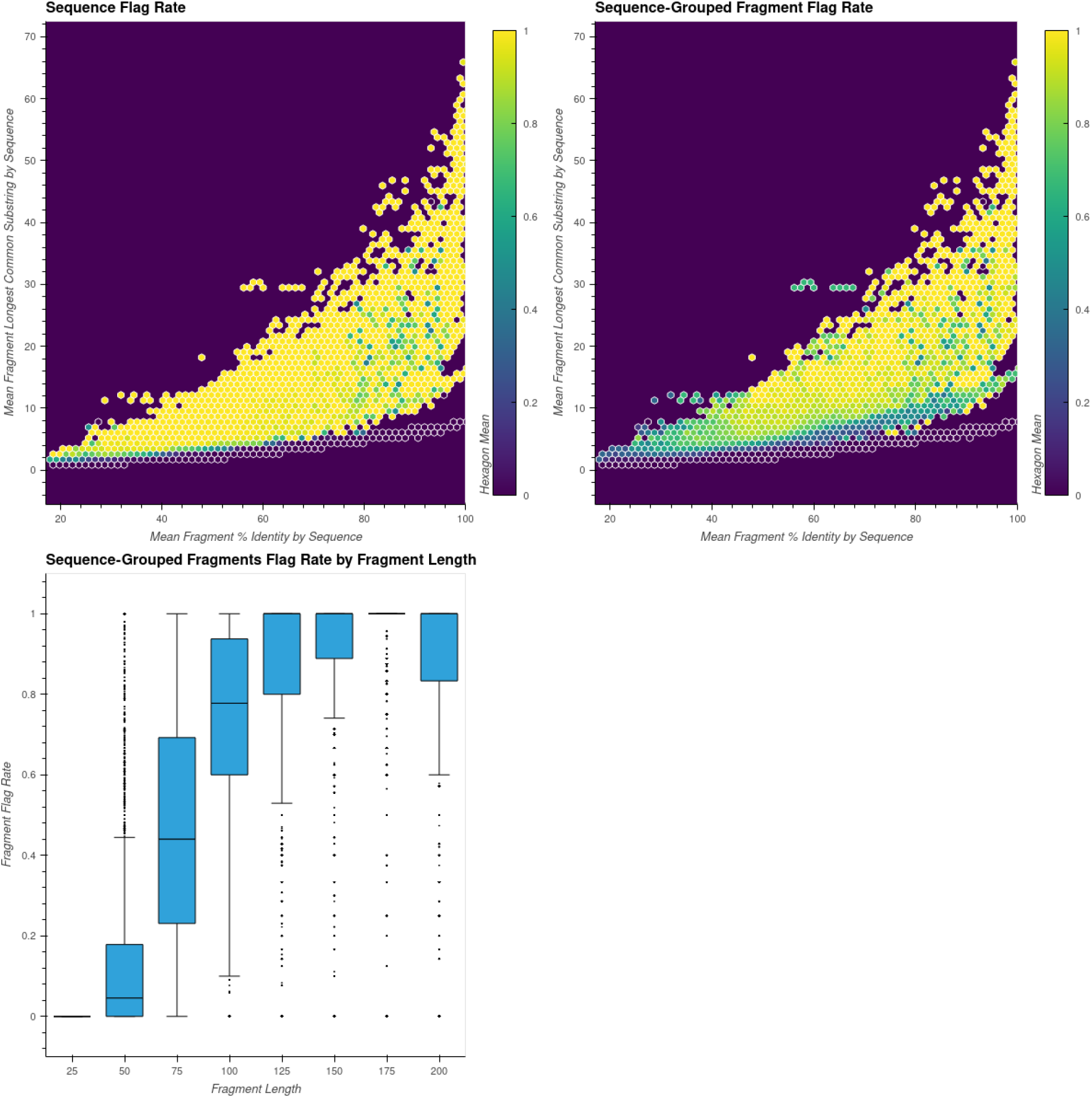
Identical to Fig. S1, but for Provider 3, Tool B.

**Fig. S5.**
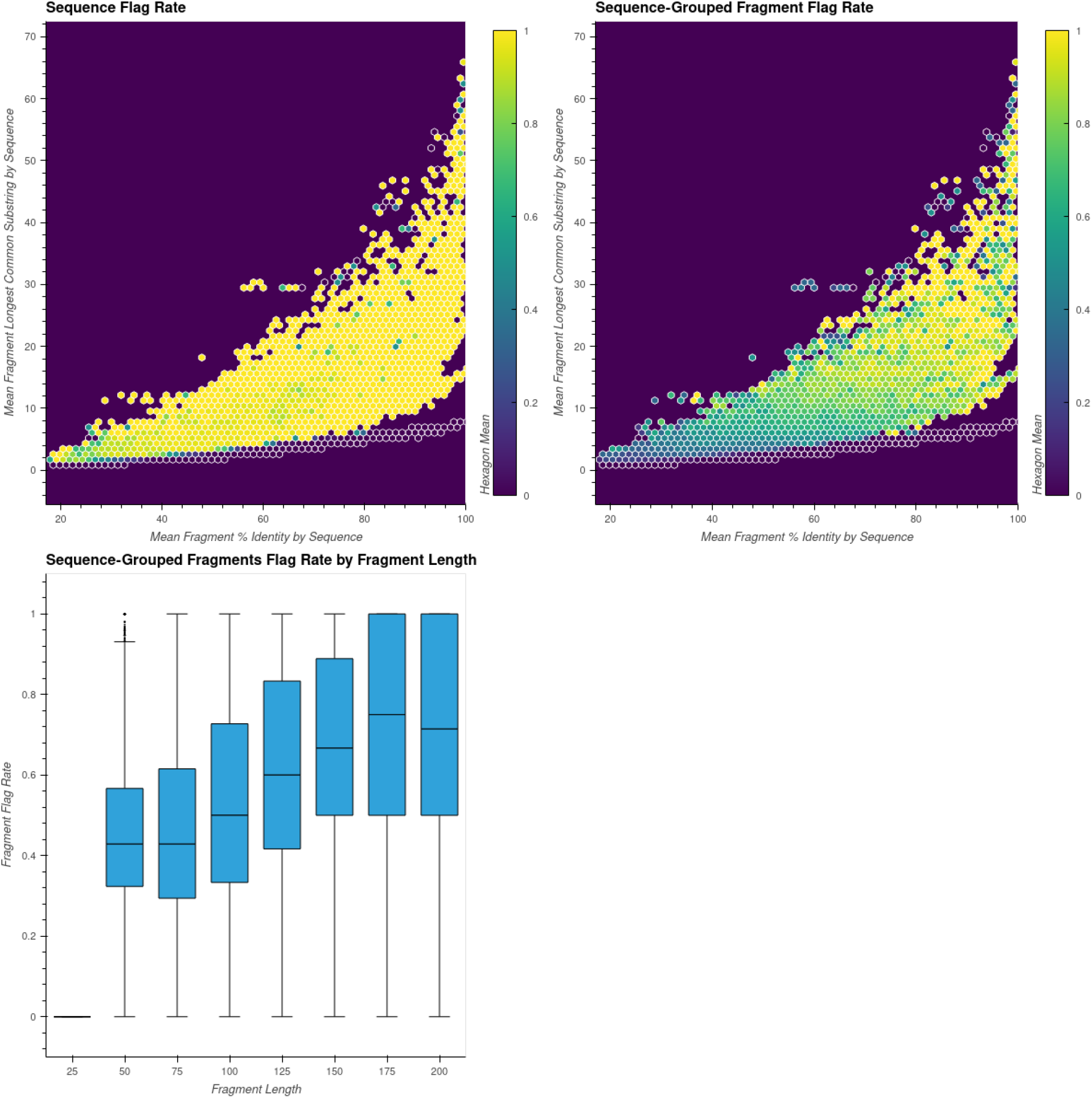
Identical to Fig. S1, but for Provider 4, Tool A.

**Fig. S6.**
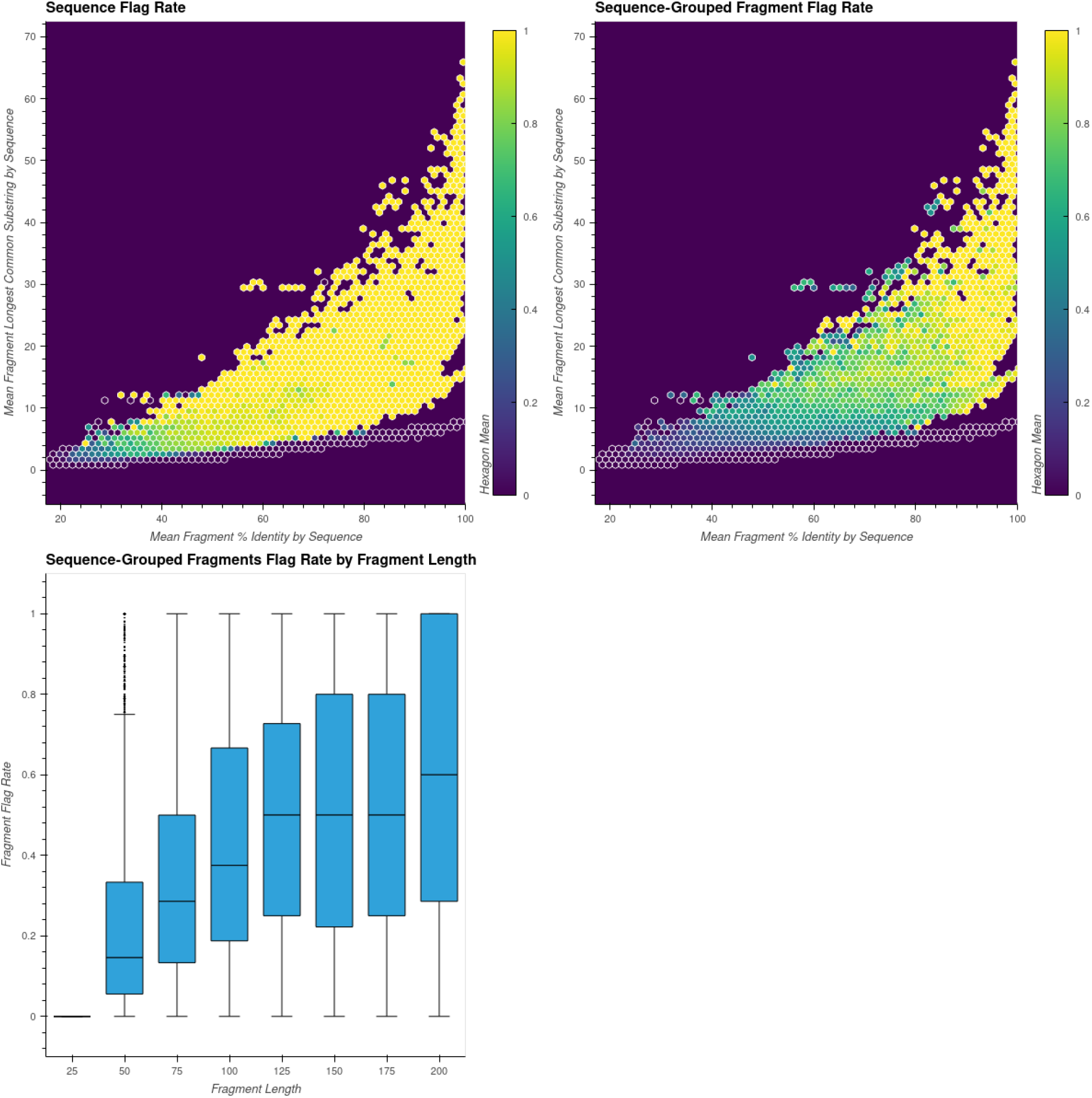
Identical to Fig. S1, but for Provider 4, Tool B.

